# Expanded diversity of novel hemoplasmas in rare and undersampled Neotropical bats

**DOI:** 10.1101/2023.06.05.543748

**Authors:** Dmitriy V. Volokhov, Lauren R. Lock, Kristin E. Dyer, Isabella K. DeAnglis, Benjamin R. Andrews, Molly C. Simonis, Sebastian Stockmaier, Gerald G. Carter, Cynthia J. Downs, M. Brock Fenton, Nancy B. Simmons, Daniel J. Becker

## Abstract

Hemotropic mycoplasmas are emerging as a model system for studying bacterial pathogens in bats, but taxonomic coverage of sampled host species remains biased. We leveraged a long-term field study in Belize to uncover novel hemoplasma diversity in bats by analyzing 80 samples from 19 species, most of which are infrequently encountered. PCR targeting the partial 16S rRNA gene found 41% of bats positive for hemoplasmas. Phylogenetic analyses found two novel host shifts of hemoplasmas, four entirely new hemoplasma genotypes, and the first hemoplasma detections in four bat species. One of these novel hemoplasmas (from *Neoeptesicus furinalis*) shared 97.6% identity in the partial 16S rRNA gene to a human hemoplasma (*Candidatus* Mycoplasma haemohominis). Additional analysis of the partial 23S rRNA gene allowed us to also designate two novel hemoplasma species, in *Myotis elegans* and *Phyllostomus discolor*, with the proposed names *Candidatus* Mycoplasma haematomyotis sp. nov. and *Candidatus* Mycoplasma haematophyllostomi sp. nov., respectively. Our analyses show that additional hemoplasma diversity in bats can be uncovered by targeting rare or undersampled host species.

## Main text

Several decades of research have characterized the diversity of viruses in bats, for which some species host viruses of notable human health concern [1]. Although bats remain relatively understudied for other pathogens, the diversity of their bacterial pathogens has been of increasing interest [2]. Hemotropic mycoplasmas (hemoplasmas) are facultative intracellular bacteria that are emerging as a model group of bat pathogens [3]. Sampling has uncovered substantial genetic diversity of bat hemoplasmas [4–6], including high similarity to a human hemoplasma (i.e., *Candidatus* Mycoplasma haemohominis, *C*Mhh) that may indicate zoonotic transmission [7,8]. However, sampling has focused on particular geographies and clades of bats, leaving taxonomic gaps in understanding how hemoplasma diversity is distributed globally and which bats may harbor potentially zoonotic hemoplasmas [2,3]. For example, most bat hemoplasma surveys have focused on species in Central and South America, with limited attention to bats in North America, Africa, Europe, and Asia [5,9]. Even within well-sampled regions of Central and South America, there is substantial variation in survey effort, with hemoplasmas of common vampire bats (*Desmodus rotundus*) being notably well-characterized while those of rarer species (e.g., *Myotis* spp., *Micronycteris* spp., *Noctilio* spp., phyllostomines such as *Mimon cozumelae* and *Trachops cirrhosus*) have been minimally sampled [4,10–13]. Here, we leveraged ongoing field studies in Belize from which we previously identified 24 new hemoplasma genotypes in 23 bat species to specifically test a range of undersampled bat species for these bacteria [4,10].

During April–May 2019, November 2021, and April–May 2022, we sampled a diverse bat community in Orange Walk District, Belize [14]. Bats were captured using mist nets and harp traps along flight paths and at roosts, held in individual bags, and identified to species based on morphology [15]. Blood volumes under 1% body mass were sampled by lancing the propatagial vein using sterile needles and collected with heparinized capillary tubes. Blood was preserved on Whatman FTA cards and held at room temperature until permanent storage at -20C at the University of Oklahoma. Field methods were approved by the Institutional Animal Care and Use Committees of the American Museum of Natural History (AMNHIACUC-20190129 and -20210614) and University of Oklahoma (2022-0197). Bat sampling was authorized by the Belize Forest Department by permits FD/WL/1/19(06), FD/WL/1/19(09), and FD/WL/1/21(12).

We used Qiagen QIAamp DNA Investigator Kits to extract genomic DNA from 80 blood samples collected from 19 bat species spanning these three years (Table 1). Our prior studies and those of other groups had mostly minimally sampled 16 of these bat species [4,6,13], whereas *Lophostoma nicaraguae, Micronycteris microtis*, and *Molossus alvarezi* remained as-of-yet wholly unsampled. To determine the presence of hemoplasmas, we used PCR targeting the partial 16S rRNA gene with primers described previously [4,10]. We also attempted to amplify the partial 23S rRNA gene for 16S rRNA–positive samples to establish a broader library of this gene for hemoplasmas [16]. Amplicons were purified and directly sequenced at Psomagen. We then used NCBI BLASTn to identify related 16S rRNA sequences, followed by MUSCLE for sequence alignment and MrBayes for phylogenetic analysis (10,000,000 generations with a GTR+G+I model) using the NGPhylogeny.fr platform [17]. We delineated hemoplasma genotypes based on analysis of the partial 16S rRNA gene (850–860 bp) and clustering on the hemoplasma phylogeny, using our previously established criteria for novelty [4,10].

**Table 1.**
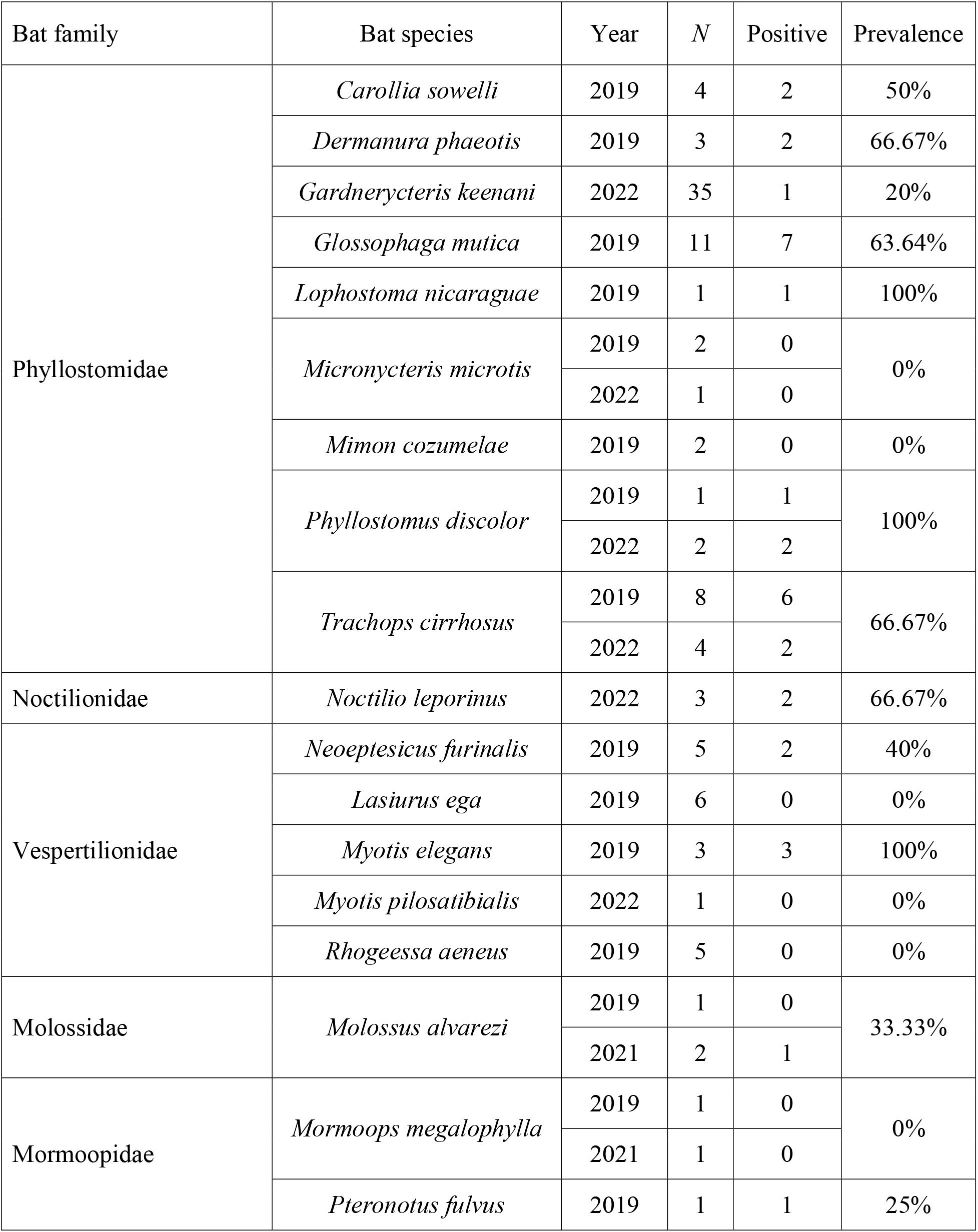
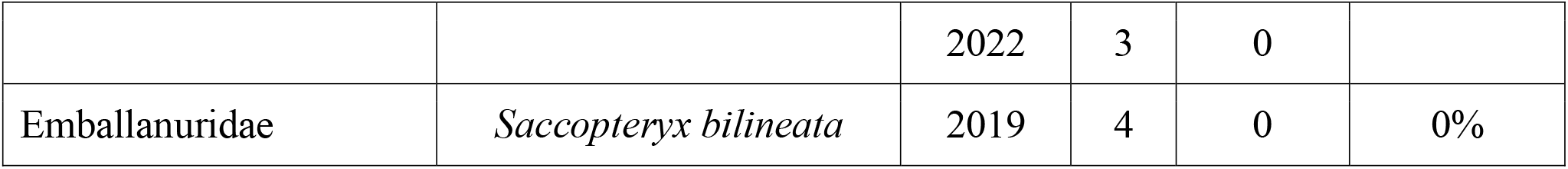
Bat samples tested for hemoplasmas in Belize, stratified by year and including 16S rRNA gene PCR positivity. We include species-level positivity for comparison to prior surveys.

We detected hemoplasmas via amplification of the partial 16S rRNA gene in 41% of samples (95% CI = 31–52%; Table 2 and Figure 1). Of the 12 bat species that tested positive, five had identical genotypes to those we earlier described in Belize [4]: *Myotis elegans* and the MYE genotype, *Carollia sowelli* and the CS1 and CS2 genotypes, *Glossophaga mutica* (formerly *G. soricina*) and the GLS genotype, *Trachops cirrhosus* and the TC1 and TC2 genotypes, and *Pteronotus fulvus* and the PPM1 genotype. All three Belize *Phyllostomus discolor* had sequences 100% identical to partial sequences from *P. discolor* in Brazil [6] and to those sequences our team has identified from *P. discolor* in Panama (GenBank Accessions OR056003–5, OR435279–81); we here classify these hemoplasmas as the PD1 genotype. For one of the PCR-positive Belize *P. discolor*, we also successfully amplified the partial 23S rRNA gene (OR055988). Owing to 100% inter-sequence identity of this sequence our recent hemoplasma 23S rRNA sequences from *P. discolor* in Panama (OR055989–92, OR435277), as well as 100% identity of the partial 16S rRNA sequences amplified from the same Belize and Panama individuals (Table 2), we propose the name *Candidatus* Mycoplasma haematophyllostomi sp. nov. to designate this novel hemoplasma species. We also amplified partial 23S rRNA genes for all three PCR-positive *Myotis elegans* (OQ456384–86). Given the high inter-sequence identity of these sequences (x□ = 99.98%) and 100% identity of paired partial 16S rRNA sequences (Table 2), we propose the name *C*. M. haematomyotis sp. nov. to designate this hemoplasma species. We also amplified the partial 23S rRNA gene for four more 16S rRNA–positive bats: *Neoeptesicus furinalis* (OQ456392), *Noctilio leporinus* (OR055987), and *Trachops cirrhosus* (OR398672–73). We were unable to establish other *Candidatus* species, as these partial 23S rRNA sequences were amplified from only one individual per bat thus far.

**Table 2.**
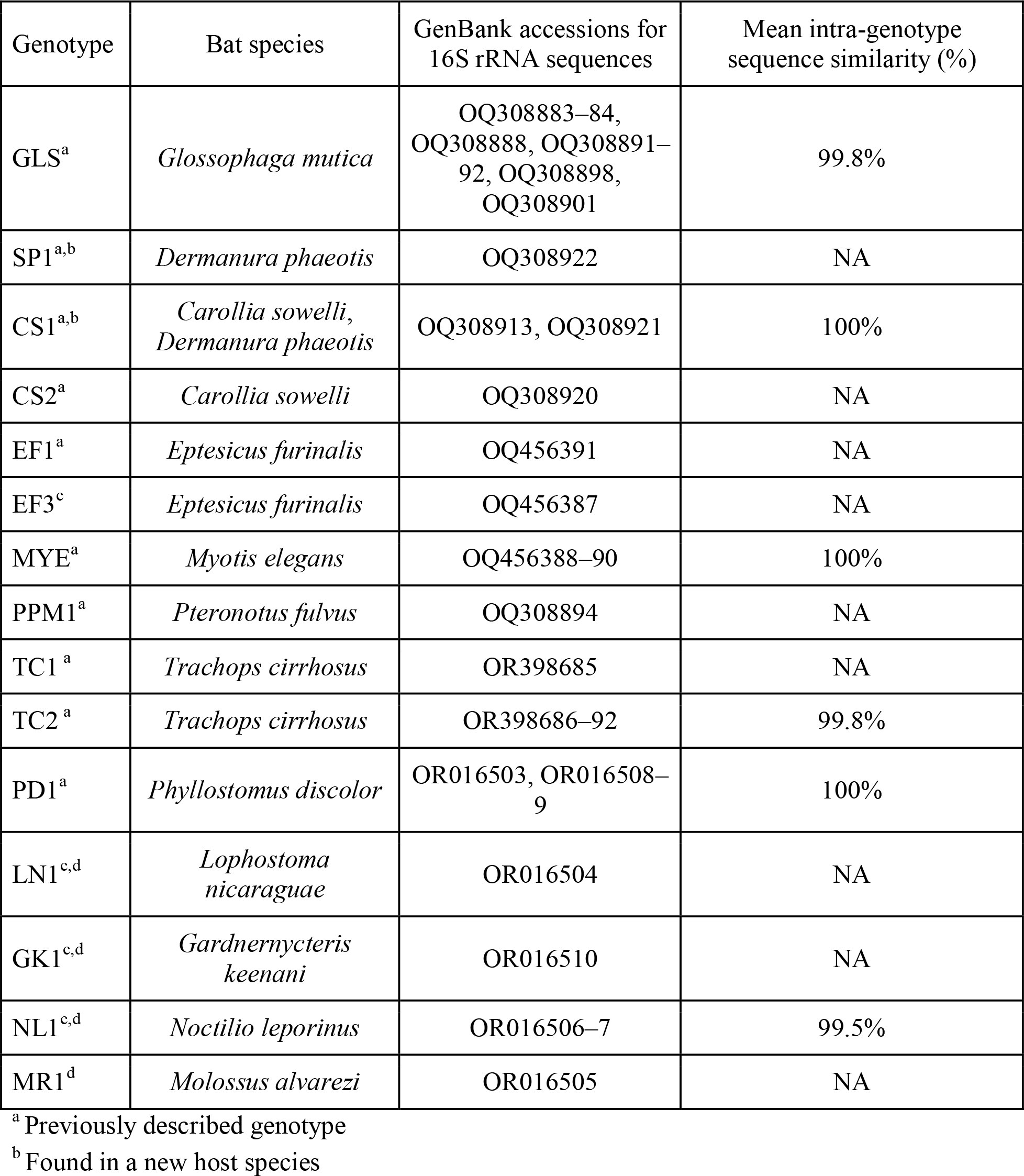
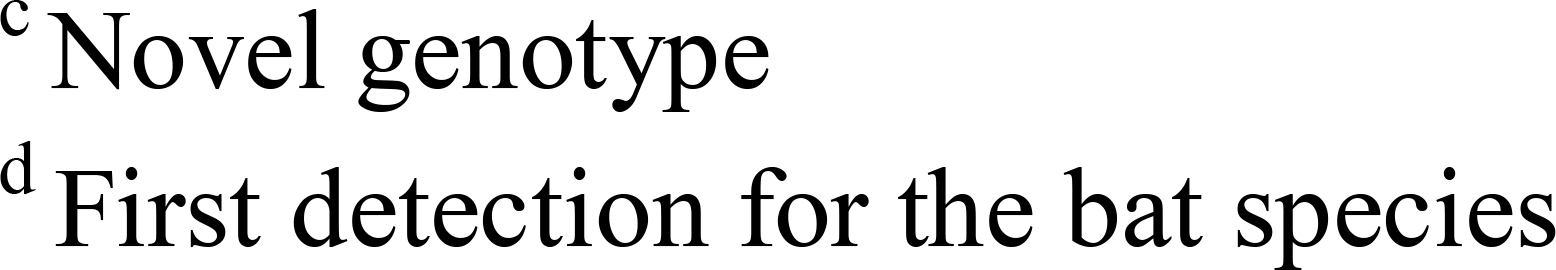
Hemoplasma genotypes identified from Belize bats. Genotypes are given with their bat host species, GenBank accessions, and intra-genotype variability from the partial 16S rRNA sequences identified in this study.

**Figure 1.**
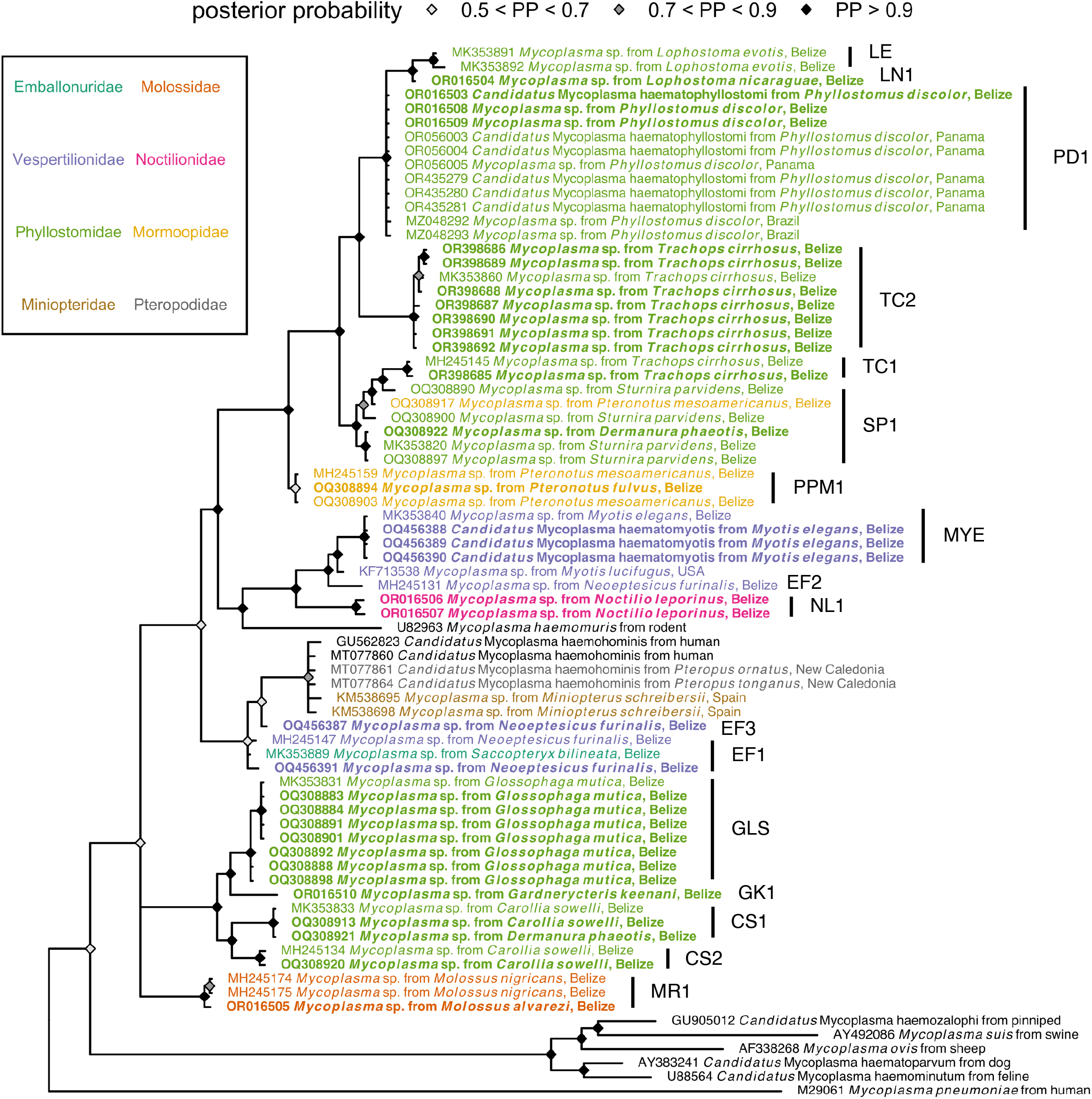
Consensus Bayesian phylogeny of the partial 16S rRNA hemoplasma sequences from this study (highlighted in bold; see Table 2 for genotype assignments) and reference sequences from bats and other mammals. Nodes are colored by posterior probability (nodes with less than 50% support are not shown), and hemoplasma sequences from bats are colored by bat family.

The remaining six bat species were infected with novel genotypes in that host (i.e., host shifts or a new host-specific genotype) or represented the first hemoplasma detections for that species. For the former, we detected two genotypes in *Dermanura phaeotis* that were previously identified as specific to *Carollia sowelli* (CS1) and *Sturnira parvidens* (SP1, group B); the later genotype was also previously detected once in *Artibeus lituratus* [4], and we recently identified the group A variant in *Pteronotus mesoamericanus* [18]. Whereas the former four bat species belong to the family Phyllostomidae, *Dermanura* and *Carollia* belong to different subfamilies in that clade, suggesting inter-generic host shifts restricted by broader evolutionary history [4].

Novel hemoplasmas were detected in *Neoeptesicus furinalis, Molossus alvarezi, Lophostoma nicaraguae, Gardnerycteris keenani*, and *Noctilio leporinus*, with the latter four representing the first hemoplasma detections in these species and the latter three representing entirely novel hemoplasma genotypes. Although we previously characterized two genotypes in *Neoeptesicus furinalis* (EF1 and EF2, with the former also identified here) [4], we identified a third, novel genotype (EF3) in a single *Neoeptesicus furinalis*, with 99.4% identity to EF1. This new EF3 genotype also was 97.6% identical to *C*Mhh isolates (e.g., MT077859 and GU562823), exceeding similarity to this human hemoplasma previously observed for the EF1 and EF2 genotypes and representing similar or greater levels of relatedness to *C*Mhh as previously observed in *Miniopterus* and *Myotis* bats in Europe and Asia as well as *Eidolon, Chaerophon*, and *Rousettus* bats in Africa [5,7,9]. However, given that the average substitution rate for the 16S rRNA gene is estimated to be approximately 1–2% per 50 million years [19], these bat hemoplasmas and *C*Mhh likely separated from a common ancestor 60–120 million years ago.

For the other bat species with novel hemoplasmas, a single *Molossus alvarezi* tested positive for the MR1 genotype (99.8% identity), which was previously identified only in the closely related *M. nigricans* in Belize [4]. We identified the novel NL1 genotype in *Noctilio leporinus*, one of two members of the family Noctilionidae. This genotype shared only 96.1% identity to the MYE genotype (i.e., *C*. M. haematomyotis sp. nov.) found in *Myotis elegans*, a distantly related vespertilionid [4]. We identified the novel LN1 genotype in one *Lophostoma nicaraguae*, which shared 99.2% identity to the LE genotype previously found in the closely related *L. evotis* [4] and 99.1% identity to the PD1 genotype (i.e., *C*. M. haematophyllostomi sp. nov.) in *P. discolor* [6]; all three bat species are members of the Phyllostominae subfamily. We lastly detected a novel genotype (GK1) in a single other phyllostomine (*Gardnerycteris keenani*), but GK1 was most related to the GLS genotype (97.6%) found prior in *Glossophaga mutica* [4].

Our analyses demonstrate additional genetic diversity of hemoplasmas in bats uncovered by targeting less-sampled species, including a novel hemoplasma with moderate phylogenetic similarity to *C*Mhh. We encourage further analyses of hemoplasmas in other unsampled bat taxa and geographies to improve our global understanding of these bacterial pathogens [2,3]. Further, longitudinal studies are needed to understand how these novel hemoplasmas circulate in bats, transmission risk to other species, and impacts on bats. Novel computational tools could also be leveraged to predict whether these new bat hemoplasmas may pose human health risks [20].

## Data availability

Hemoplasma sequence data are available on GenBank for the partial 16S rRNA gene (Table 2) and partial 23S rRNA gene (accessions OR055988, OQ456384–86, OQ456392, OR055987, and OR398672–73). PCR positivity data are summarized in Table 1 and are fully available in the Pathogen Harmonized Surveillance (PHAROS) database: https://pharos.viralemergence.org/.

## Acknowledgements

This work was supported by the National Geographic Society (NGS-55503R-19), SUNY-ESF Honors Program, Explorer’s Club, and Research Corporation for Science Advancement (RCSA). This work was conducted as part of Subaward No. 28365, part of a USDA Non-Assistance Cooperative Agreement with RCSA Federal Award No. 58-3022-0-005. MCS was supported by an appointment to the Intelligence Community Postdoctoral Research Fellowship Program at University of Oklahoma administered by Oak Ridge Institute for Science and Education through an interagency agreement between the U.S. Department of Energy and the Office of the Director of National Intelligence. SS was supported by an Ohio State University President’s Postdoctoral Fellowship. GGC and DJB were supported by the National Science Foundation (IOS 2015928 and BII 2213854, respectively). We thank Mark Howells and staff of the Lamanai Field Research Center for assistance with field logistics and permits as well as the many colleagues who helped net bats during 2019, 2021, and 2022 research in Belize. We thank Konstantin Chumakov for laboratory support and two reviewers for feedback on a previous version of this manuscript.

